# Different RNA profiles in plasma derived small and large extracellular vesicles of Neurodegenerative diseases patients

**DOI:** 10.1101/2020.11.23.390591

**Authors:** Daisy Sproviero, Stella Gagliardi, Susanna Zucca, Maddalena Arigoni, Marta Giannini, Maria Garofalo, Martina Olivero, Michela Dell’Orco, Orietta Pansarasa, Stefano Bernuzzi, Micol Avenali, Matteo Cotta Ramusino, Luca Diamanti, Brigida Minafra, Giulia Perini, Roberta Zangaglia, Alfredo Costa, Mauro Ceroni, Nora I. Perrone-Bizzozero, Raffaele A. Calogero, Cristina Cereda

## Abstract

**Background:** Identifying robust biomarkers is essential for early diagnosis of neurodegenerative diseases (NDs). Large (LEVs) and small extracellular vesicles (SEVs) are extracellular vesicles (EVs) of different sizes and biological functions transported in blood and they may be valid biomarkers for NDs. The aim of our study was to investigate common and different mRNA/miRNA signatures in plasma derived LEVs and SEVs of Alzheimer’s Disease (AD), Parkinson’s disease (PD), Amyotrophic Lateral Sclerosis (ALS) and Fronto-Temporal Dementia (FTD) patients.

**Methods:** LEVs and SEVs were isolated from plasma of patients and healthy volunteers (CTR) by filtration and ultracentrifugation and RNA was extracted. Whole transcriptome and miRNA libraries were carried out by Next Generation Sequencing (NGS).

**Results:** We detected different deregulated RNAs in LEVs and SEVs from patients with the same disease. MiRNAs resulted to be the most interesting subpopulation of transcripts transported by plasma derived SEVs since they appeared to discriminate all NDs disease from CTRs and they can provide a signature for each NDs. Common enriched pathways for SEVs were mainly linked to ubiquitin mediated proteolysis and Toll-like receptor signaling pathways and for LEVs to neurotrophin signaling and Glycosphingolipid biosynthesis pathway.

**Conclusion:** LEVs and SEVs are involved in different pathways and this might give a specificity to their role in the spreading/protection of the disease. The study of common and different RNAs transported by LEVs and SEVs can be of great interest for biomarker discovery and for pathogenesis studies in neurodegeneration.

## Background

Neurodegenerative disorders are a group of diseases characterized by loss of neurons within the brain and/or spinal cord [1]. They include both common neurodegenerative disorders such as Alzheimer’s disease (AD), Parkinson’s disease (PD), and rare diseases as Amyotrophic Lateral Sclerosis (ALS) and Fronto-Temporal Dementia (FTD) [2]. Each of these disorders are characterized by specific features, both clinical and pathological involving characteristic central nervous regions [3].

Genetics are also known to play an essential role in neurodegenerative diseases (NDs). Genome-wide association studies (GWAS) identified many causative mutations and single nucleotide polymorphisms (SNPs) in genes associated with the etiology or risk factors of NDs. However, only few cases of NDs can be explained by a typical Mendelian inheritance and more than 90% of cases, defined as “sporadic” forms, are regulated by other pathways [4].

Neurodegenerative diseases have in common protein aggregates such as pathological hallmark lesions. Amyloid β-protein, Tau-protein, α-synuclein, TDP-43 and SOD1 are the most frequently aggregated proteins [5]. AD is characterized by the extracellular accumulation of beta amyloid (Aβ) peptide detectable as Aβ plaques in patients’ brain. Also intracellular Tau protein aggregation is found in AD brain [6]. The main PD features are degeneration of the dopaminergic pigmented neurons in the substantia nigra (SN) and accumulation of α-synuclein protein, which is the main component of Lewy bodies (LBs) [7]. ALS is a disease characterized by motor neurons death and one of the main pathological hallmarks is given by specific alterations of SOD1 [8,9,10], and aggregation of TDP-43 [11]. In FTD, there is a deregulation of RNA-binding proteins (RBPs) and aggregation of proteins in the frontal and temporal lobes with microvacuolation, neuronal loss and astrocytic gliosis [12]. TDP-43 and Tau aggregates are hallmarks of FTD [13]. Although these proteins constitute disease-characteristic aggregates, they are not restricted to these clinical presentations [5]. For example, TDP-43 pathology is also found in over 50% of cases of AD patients, while Fronto-Temporal Lobar Degeneration (FTLD) can be subclassified in disorders that accumulate τ, i.e., FTLD-tau; TDP-43, i.e., FTLD-TDP, and other FTLD-forms that accumulate other proteins (e.g., fused in sarcoma (FUS) [14]. FTLD-TDP and ALS share neuronal TDP-43 aggregates and a certain number of ALS cases also develop FTLD-TDP [11,15]. In NDs, the abnormal protein accumulation triggers the activation of common biological and molecular features, inflammatory and oxidative-stress pathways and mitochondrial dysfunction [16,17,18,19,20].

Although the role of protein aggregates received much emphasis in NDs, the aberrant RNA metabolism processing converges as a common factor in the pathogenesis of these diseases [21,22]. Abnormal RNA metabolism is associated with disease-specific alterations in RNA-binding proteins (RBPs), and in non-coding RNAs, such as microRNAs (miRNA), transfer RNAs (tRNA) and long-noncoding RNAs (lncRNA) [23].

Several common and specific mechanisms of NDs are described in the literature, however no biomarkers are available to identify the onset, progression and comorbidity of those diseases. Brain cells release extracellular vesicles (EVs), which can pass brain barrier [24,25] and blood derived EVs can be used also to monitor disease processes occurring in the brain [26,27,28]. EVs, spherical vesicles heterogeneous in size (30 nm-1 μm in diameter) are transporters of receptors, bioactive lipids, proteins, and nucleic acids, such as mRNAs, lncRNAs and miRNAs [24,25,26,27]. EVs are classified as: exosomes (EXOs), microvesicles (MVs), and apoptotic bodies [29,30]. EXOs are secreted membrane vesicles (approximately 30–150 nm in diameter) formed intracellularly and released from exocytosis of multivesicular bodies, whereas apoptotic bodies (approximately 1000–4000 nm in diameter) are released by dying cells. MVs (approximately 100–1000 nm in diameter) are shed from cells by outward protrusion (or budding) of a plasma membrane followed by fission of their membrane stalk [29,30]. However, the guidelines of the International Society for the study of Extracellular Vesicle (ISEV) released in 2018, declare that MVs and EXOs cannot be distinguished on a particular biogenesis pathway and so they can be distinguished in small extracellular vesicles (SEVs) (30-130 nm) and large extracellular vesicles (LEVs) (130-1000 nm) mainly on their size [31].

Several studies on the role of extracellular vesicles in NDs are available in the literature. Some studies have examined miRNAs and RNAs in EVs isolated from cultured cell media from the Central Nervous System (CNS) cells (e.g., neurons, astrocytes, microglia, and oligodendrocytes) and few have examined miRNAs in EVs in plasma of AD, PD and ALS [32, 33, 34]. However, none of these studies considered the differences between SEVs from LEVs. We previously described in ALS that SEVs and LEVs in plasma are different in dimensions and for loading of some pathological proteins for ALS (SOD1, TDP-43, p-TDP-43, and FUS) and lipids [35,55]. In this paper, we have investigated the RNA cargo of EVs derived from plasma of patients affected by four neurodegenerative diseases (AD, PD, ALS and FTD). The aim was to identify common and specific transcripts and small RNAs s between the two subpopulation of EVs in the same disease and in the four diseases in order to identify new biomarkers.

## Methods

### Study Subjects

Participants were recruited at the IRCCS Mondino Foundation, Pavia (Italy). Subjects participating in the study signed, before being enrolled, an informed consent form approved by the Ethical Committee (for ALS patients Protocol n°-20180034329; for PD patients Protocol n°20170001758; for AD patients Protocol n°20170016071; for FTD patients Protocol n°20180049077). Plasma isolated from 6 AD, 9 PD, 6 sporadic ALS (SALS), 9 FTD patients were used (Table 1). All patients were screened for mutations using a customized panel of 176 genes associated to neurodegenerative and neuromuscular diseases by Next Generation Sequencing (Sure Select QXT Target Enrichment, Agilent Technology).

**Table 1.** Baseline characteristics of recruited subjects for this study. Age is reported as mean ± SD. The percentage of male and female subjects is also indicated.

|  | CTRs | AD | FTD | ALS | PD |
| --- | --- | --- | --- | --- | --- |
| **Recruited subjects** | 6 | 6 | 9 | 6 | 9 |
| **Age (mean±SD)** | 55±5,2 | 77±3,7 | 60±6,7 | 72±6,3 | 69±3,6 |
| **Males %** | 43% | 50% | 78% | 50% | 50% |
| **Females %** | 67% | 50% | 22% | 50% | 50% |

CTRs= controls; AD= Alzheimer Disease; FTD= Fronto-Temporal Dementia; ALS= Amyotrophic Lateral Sclerosis; PD= Parkinson Disease; SD= standard deviation.

ALS and FTD patients were screened for C9orf72 using the FastStart Taq DNA Polymerase Kit (Roche).

Diagnosis of AD was based on criteria expressed by Aging-Alzheimer’s Association workgroups [36]. For PD and FTD patients Movement Disorder Society (MDS) clinical diagnostic criteria were used [37, 38]. ALS diagnosis was made according to the revised El Escorial Criteria [39].

Six age-matched healthy volunteers free from any pharmacological treatment were recruited at the Immunohematological and Transfusional Service IRCCS Foundation “San Matteo”, Pavia (Italy) and used as healthy controls (CTRs). All the subjects were assayed to rule out the presence of inflammatory diseases by white blood cell counts and subjects with WBCs >11×10^9^ were excluded from the study. Patients’ characteristics are reported in Table 2.

**Table 2.**
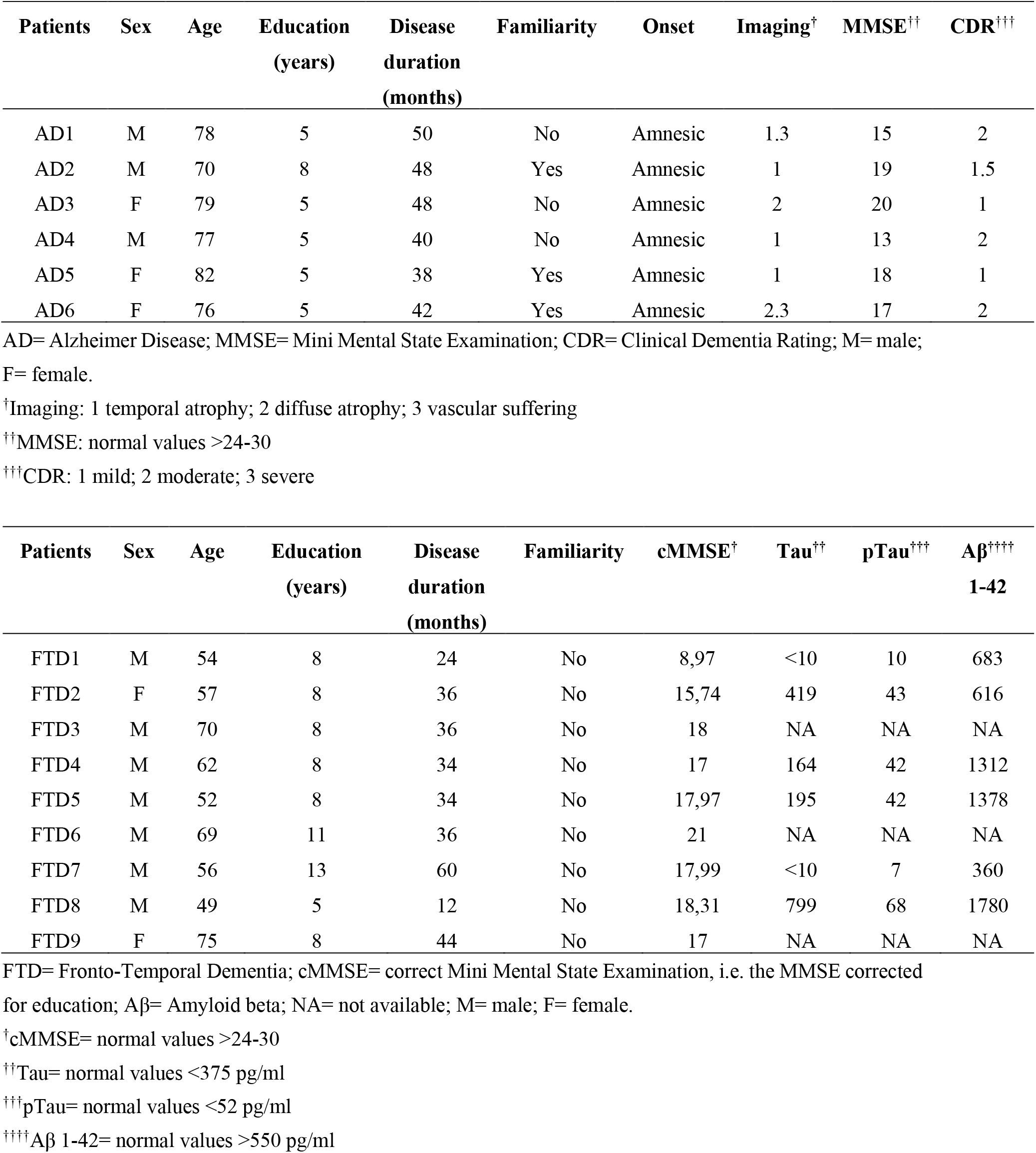
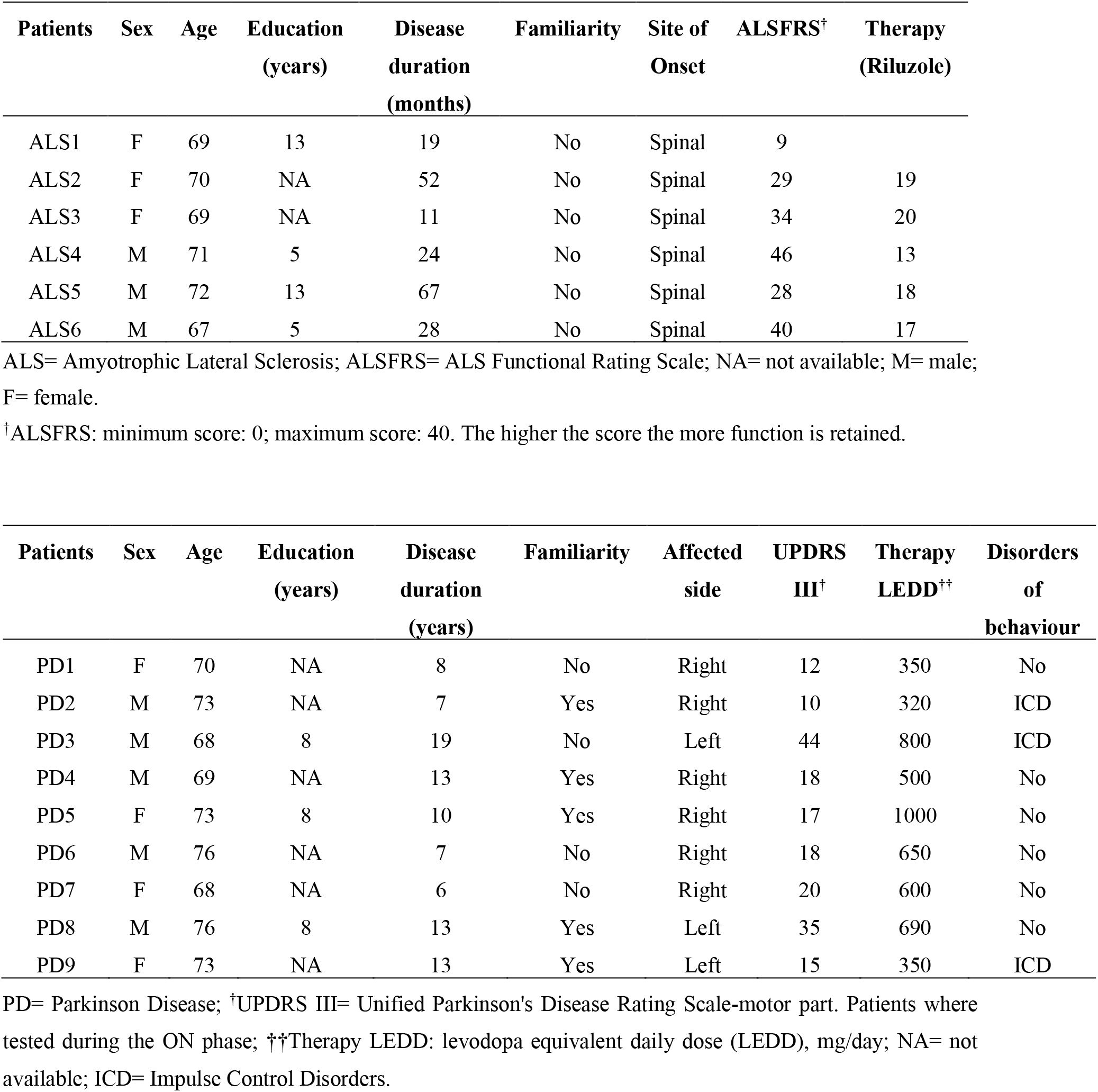
Clinical characteristics of the recruited subjects.

### LEVs and SEVs isolation

Venous blood (15 ml) was collected in sodium citrate tubes from all patients and controls and processed as previously described [35,40]. Briefly, platelet-free plasma was centrifuged at 20,000xg for 1 hour. The pellet was washed in 0.2 μm filter filtered 1X PBS (Sigma-Aldrich, Italy). The supernatant of LEVs was filtered through a 0.2 μm filter and spun in an Optima MAX-TL Ultracentrifuge at 100,000xg for 1 hour at 4°C and SEVS pellet was washed with 1 ml of filtered 1X PBS. Western Blot analysis for LEVs markers (Annexin V-Abcam, Inc., United States) and for SEVs markers (Alix-Abcam, Inc., United States), Transmission Electron Microscopy (TEM) and Nanoparticle-tracking analysis (NTA) were run to confirm LEVs and SEVs purity as we previously described [35,40].

### RNA extraction

RNA was extracted from LEVs and SEVs fractions using Qiagen miRNeasy Mini kit (Qiagen, Germany) according to the manufacturer’s instructions.

### RNA libraries preparations

Small RNA libraries were constructed from the RNA samples using NEBNext® kit (New England Biolabs, USA). Individually-barcoded libraries were mixed. Pools were size selected on Novex 10% TBE gels (Life Technologies, USA) to enrich for miRNAs fraction. Sequencing (75 nts single-end) was performed on Illumina NextSeq500 (Illumina, USA).

Long RNA libraries (mRNAs and lncRNAs) were prepared with the Illumina TruSeq Stranded RNA Library Prep (Illumina, USA). Sequencing (75 nts paired-end) was performed on Illumina NextSeq500 (Illumina, USA). Demultiplexing was done as described for miRNA sequences.

### Bioinformatic data analysis

The raw bcl files were converted into demultiplexed fastq files with bcl2fastq (Illumina, USA) implemented in docker4seq package [41]. For the row count analysis, only transcripts with counts above five were considered. No relevant difference between count in SEVs and LEVs in four diseases emerged.

Quantification of miRNAs was done as described in the literature [42]. The workflow, including quality control filter, trimming of adapters, reads mapping against miRNA precursors, is implemented in docker4seq package [42]. Differential expression analysis was performed with the R package DESeq2, implemented in docker4seq package. We imposed a minimum |Log2FC| of 1 and a FDR lower than 0.1 as thresholds to detect differentially expressed miRNAs.

Quantification of genes and isoforms was performed as previously described [43,44]. Differential expression analysis for mRNAs was performed using R package EBSeq [45], using same threshold indicated above for miRNAs differential expression analysis. In order to understand common miRNAs of the four diseases, we calculated the intersection of deregulated miRNAs compared to CTRs with http://bioinformatics.psb.ugent.be/webtools/Venn/.

The datasets generated and analysed during the current study are available in the NCBI GEO repository [GSE155700].

### Pathways analysis

Gene enrichment analysis was performed on coding genes with KEGG pathway analysis (Kyoto Encyclopedia of Genes and Genomes, http://www.genome.ad.jp/kegg) and enrichR web tool [46,47]. miRNA-targets analysis was done with miRWalk web tool (http://mirwalk.umm.uni-heidelberg.de).

In addition, Ingenuity pathway analysis (IPA, v. 2019 summer release, Qiagen, Germany) was performed to identify mRNAs and biological networks associated with the differentially expressed circulating miRNAs in SEVs and LEVs in each of the neurodegenerative disorders.

## Results

### miRNAs selectively traffic into SEVs and LEVs

We previously demonstrated significant differences between LEVs and SEVs derived from plasma for dimension, markers, protein loading (see figure S1a, b, c, d) [35,40,55]. Cellular miRNAs and RNAs can selectively traffick into LEVs and SEVs, so we first identified differentially expressed miRNAs (DE miRNAs) and mRNAs (DE mRNA) in SEVs and LEVs among the four groups of patients (AD, PD, ALS, FTD) and the healthy controls (CTRs) (Table 3, Table 4, Table S1, Table S2). We then moved to investigate the number of different and common deregulated miRNAs and RNAs that sort into SEVs and LEVs in the same disease. In AD, of the 33 miRNAs found in SEVs and 13 in LEVs, 6 distribute to both (Figure 1a), 4 upregulated and 2 downregulated. In the case of FTD, of the 88 miRNAs in SEVs and 130 in LEVs, 34 were in common, 32 upregulated and 2 downregulated (Figure 1b). Concerning miRNAs in ALS, of the 109 miRNAs in SEVs and 197 in LEVs, 67 were in common, (Figure 1c), 45 upregulated and 22 downregulated. In PD, of the 104 miRNAs found in SEVs and 109 in LEVs, 34 distribute to both (Figure 1d), 30 upregulated and 4 downregulated. For mRNA, we could only calculate the intersection between SEVs and LEVs for ALS and FTD, since there were no deregulated mRNAs (with a Log2FC| of 1) in EVs from AD patients and in SEVs of PD patients compared to CTRs. For FTD, of the 228 mRNA in SEVs and 114 in LEVs, 39 were in common, 36 upregulated and 3 downregulated (Figure 2a). For ALS, of the 522 mRNA in SEVs and 124 in LEVs, 44 were in common (33 upregulated and 11 downregulated) (Figure 2b). The percentage of common miRNAs and RNAs between SEVs and LEVs are shown in Table 5.

**Table 3.** Statistically significant number of differentially expressed miRNAs in SEVs and LEVs from ALS, FTD, AD, PD patients. Up-regulated transcripts, down-regulated transcripts and total compared to CTRs were reported. Transcripts were considered as differentially expressed when |log2(disease sample/healthy controls)|≥1 and a FDR ≤ 0.1.

|  | AD |  | FTD |  | ALS |  | PD |  |
| --- | --- | --- | --- | --- | --- | --- | --- | --- |
|  | *SEVs* | *LEVs* | *SEVs* | *LEVs* | *SEVs* | *LEVs* | *SEVs* | *LEVs* |
| **miRNA up-regulated** | 17 | 10 | 49 | 113 | 80 | 128 | 85 | 87 |
| **miRNA down-regulated** | 16 | 3 | 39 | 17 | 29 | 69 | 19 | 22 |
| **Total** | 33 | 13 | 88 | 130 | 109 | 197 | 104 | 109 |

AD= Alzheimer Disease; FTD= Fronto-Temporal Dementia; ALS= Amyotrophic Lateral Sclerosis; PD= Parkinson Disease; CTRs= controls; SEVs= small extracellular vesicles; LEVs= large extracellular vesicles; miRNA= microRNA; FDR= False Discovery Rate.

**Table 4.** Number of statistically significant differentially expressed RNAs in SEVs and LEVs from AD, FTD, ALS and PD patients in terms of up-regulated transcripts, down-regulated transcripts and total compared to CTRs. Transcripts were considered as differentially expressed when |log2(disease sample/healthy control)|≥1 and a FDR ≤ 0.1.

|  | AD |  | FTD |  | ALS |  | PD |  |
| --- | --- | --- | --- | --- | --- | --- | --- | --- |
| *WHOLE*<br>*TRANSCRIPTOME* | *SEVs* | *LEVs* | *SEVs* | *LEVs* | *SEVs* | *LEVs* | *SEVs* | *LEVs* |
| **mRNA up-regulated** | 0 | 0 | 194 | 64 | 499 | 32 | 0 | 2 |
| **mRNA down-regulated** | 0 | 0 | 33 | 23 | 43 | 56 | 0 | 10 |
| **Total** | 0 | 0 | 227 | 87 | 542 | 88 | 0 | 12 |
| **lncRNA up-regulated** | 0 | 0 | 15 | 17 | 16 | 14 | 0 | 0 |
| **lncRNA down-regulated** | 0 | 0 | 0 | 2 | 0 | 4 | 0 | 1 |
| **Total** | 0 | 0 | 15 | 19 | 16 | 18 | 0 | 1 |

AD= Alzheimer Disease; FTD= Fronto-Temporal Dementia; ALS= Amyotrophic Lateral Sclerosis; PD= Parkinson Disease; CTRs= controls; SEVs= small extracellular vesicles; LEVs= large extracellular vesicles; lncRNAs= long non coding RNAs, FDR= False Discovery Rate.

**Table 5.** Percentage of common miRNAs and mRNAs in SEVs and LEVs in AD, FTD, ALS and PD.

| miRNA |  |  |
| --- | --- | --- |
| Common miRNAs in SEVs and LEVs (%) |  |  |
|  | *SEVs* | *LEVs* |
| **AD** | 18.2 | 46.2 |
| **FTD** | 38.6 | 25.2 |
| **ALS** | 61.5 | 34 |
| **PD** | 32.7 | 31.2 |
| mRNA |  |  |
| Common mRNAs in SEVs and LEVs (%) |  |  |
|  | *SEVs* | *LEVs* |
| **AD** | 0 | 0 |
| **FTD** | 17.1 | 34.2 |
| **ALS** | 8.4 | 35.5 |
| **PD** | 0 | 0 |

AD= Alzheimer Disease; FTD= Fronto-Temporal Dementia; ALS= Amyotrophic Lateral Sclerosis; PD=
Parkinson Disease; CTRs= controls; SEVs= small extracellular vesicles; LEVs= large extracellular vesicles;
miRNAs= micro RNAs.

**Figure 1.**
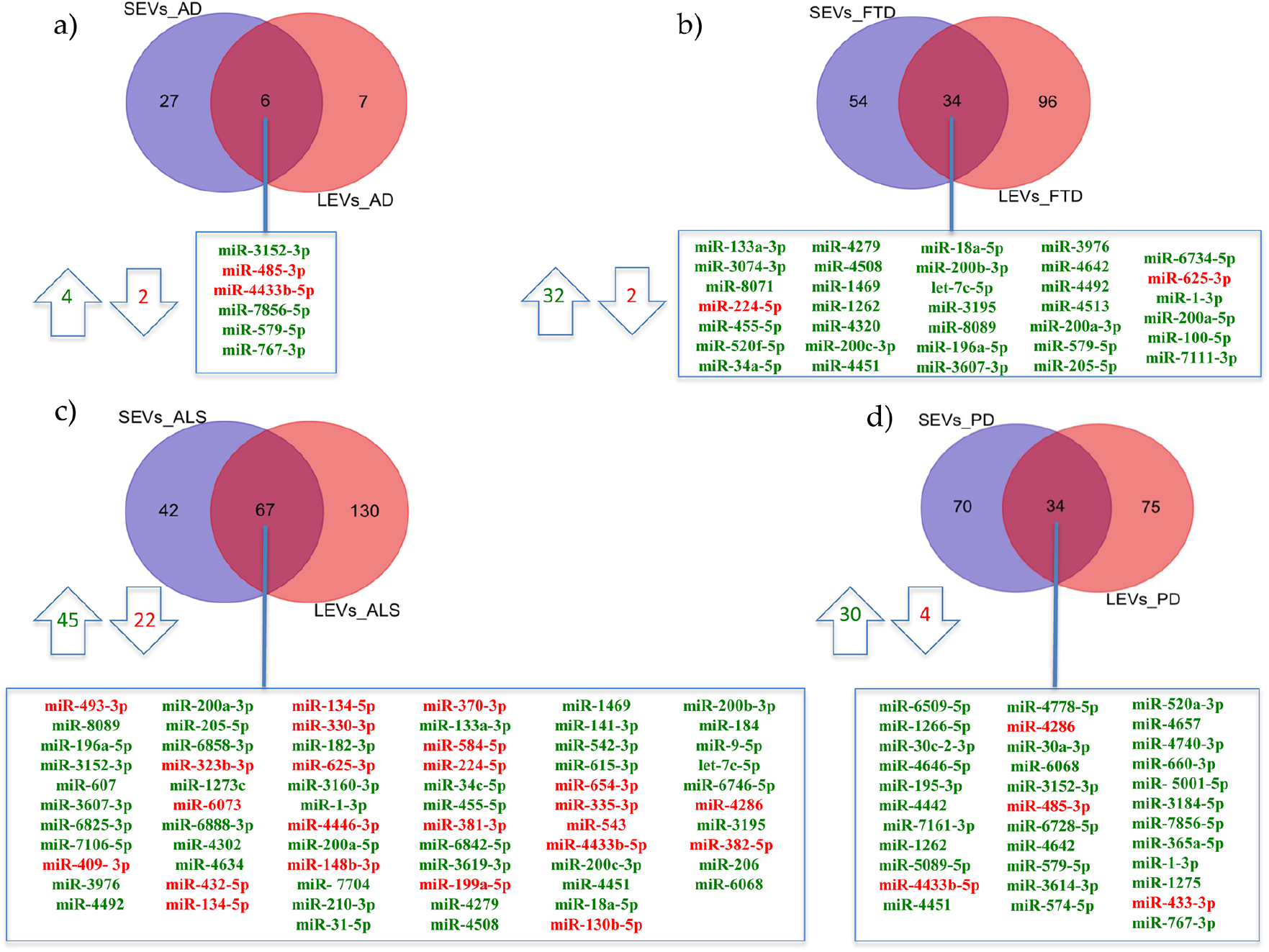
Common packaging of deregulated miRNAs into SEVs and LEVs of NDs. (a) In AD, of the 33 miRNAs found in SEVs and 13 in LEVs, 6 distribute to both, 4 upregulated-in green and 2 downregulated-in re;. (b) In FTD, of the 88 miRNAs in SEVs and 130 in LEVs, 34 were in common (32 upregulated and 2 downregulated); c) for ALS, of the 109 miRNAs in SEVs and 197 in LEVs, 67 were in common, 45 upregulated and 22 downregulated); d) in PD, of the 104 miRNAs found in SEVs and 109 in LEVs, 34 distribute to both, 30 upregulated and 4 downregulated. Differential miRNA expression analysis by DESeq2 (log2FC > 1, p-value<0.05).

**Figure 2.**
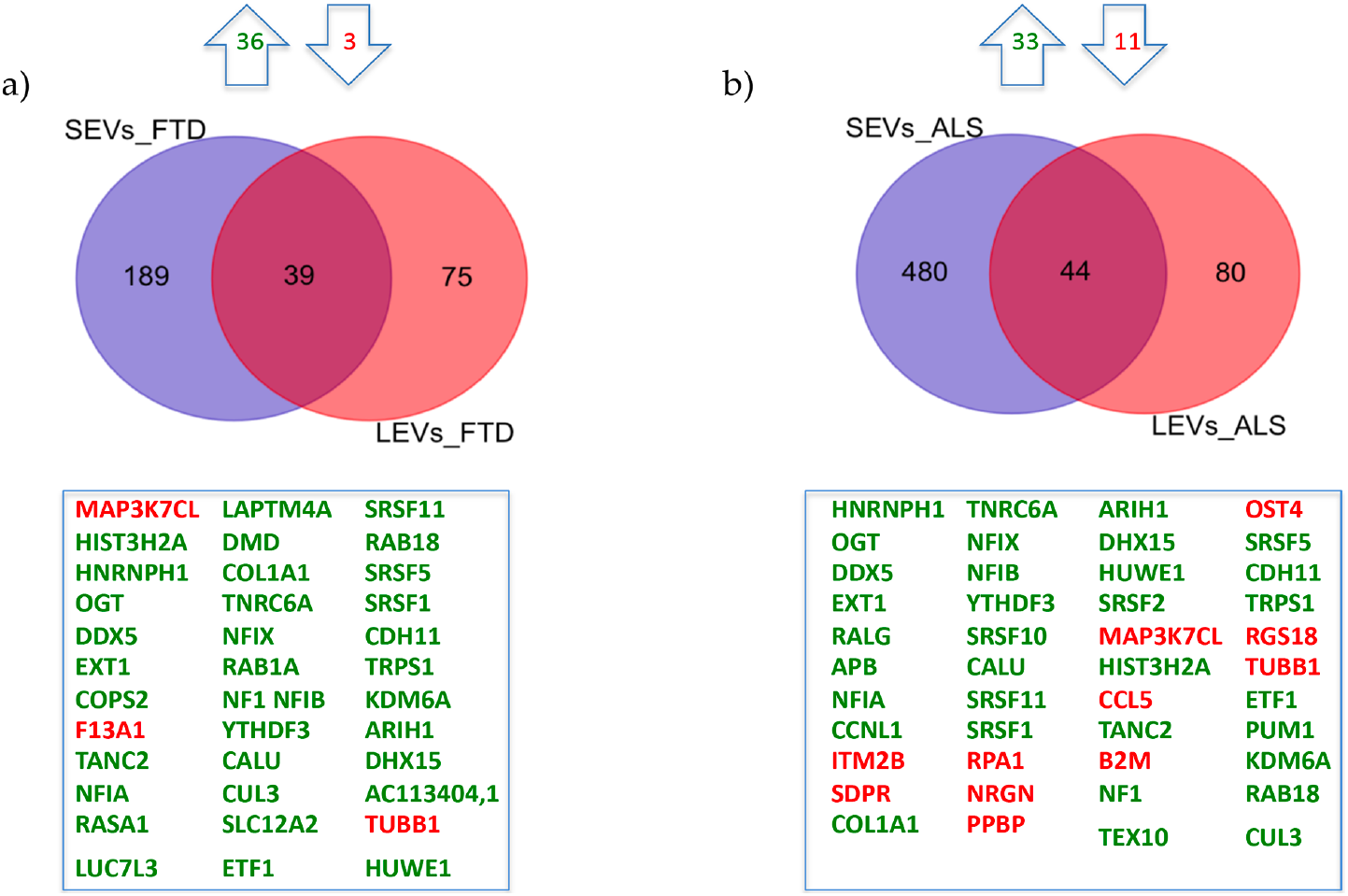
Common packaging of deregulated mRNAs into SEVs and LEVs from FTD and ALS patients. (a) For FTD, of the 228 mRNA in SEVs and 114 in LEVs, 39 were in common (36 upregulated, green and 3 downregulated, red). (b) For ALS, of the 522 mRNA in SEVs and 124 in LEVs, 44 were in common (33 upregulated and 11 downregulated). Differential mRNA expression analysis by DESeq2 (log2FC > 1, p-value<0.05).

### miRNAs expression profiles and common pathways in SEVs and LEVs of NDs

miRNAs detected as differentially expressed in SEVs at least one disease were pooled together and analyzed by principal component analysis (PCA) (Figure 3a). The same approach was applied to LEVs.

**Figure 3.**
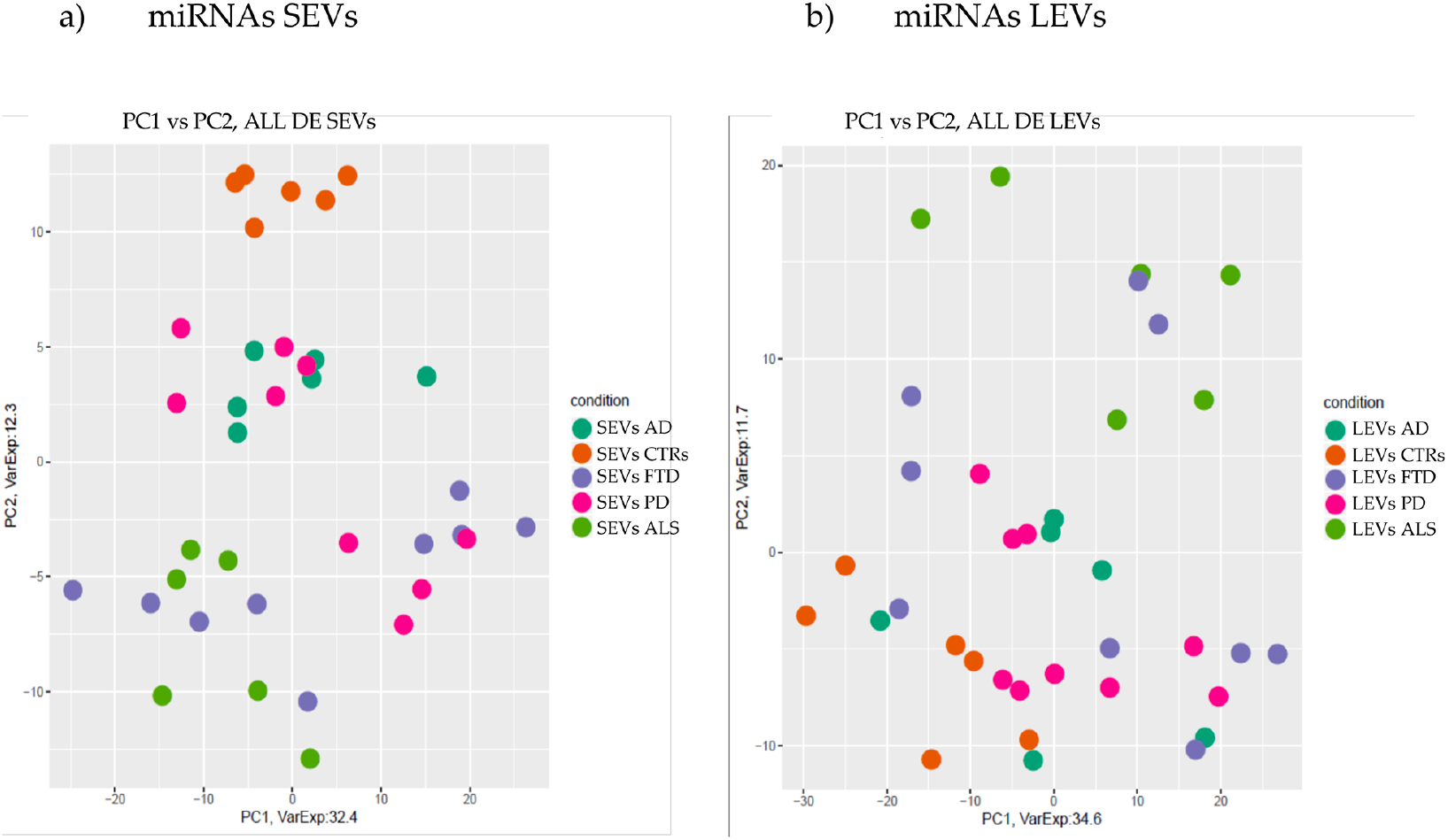
PCA of miRNAs differentially expressed in SEVs (a) and LEVs (b) of ALS, FTD, AD and PD patients and healthy controls (CTRs). PCA is performed using as predictors all the miRNAs identified as differentially expressed in at least one disease in the comparison of each disease to the control state. Each dot represents a sample and each color represents a disease.

As shown in Figure 3a, miRNAs contained in SEVs of the four NDs did not overlap with the CTRs (orange). Interestingly, clusters related to ALS (light green) and AD (dark green) patients could be identified in separate components, while (violet) overlapped with some ALS specimens. The miRNA cargo of SEVs were well differentiated in all four diseases; interestingly, we identified a well-defined group of specific miRNAs in SEVS from ALS, suggesting a possible signature for the disease (Figure 3a, Figure 4a).

**Figure 4.**
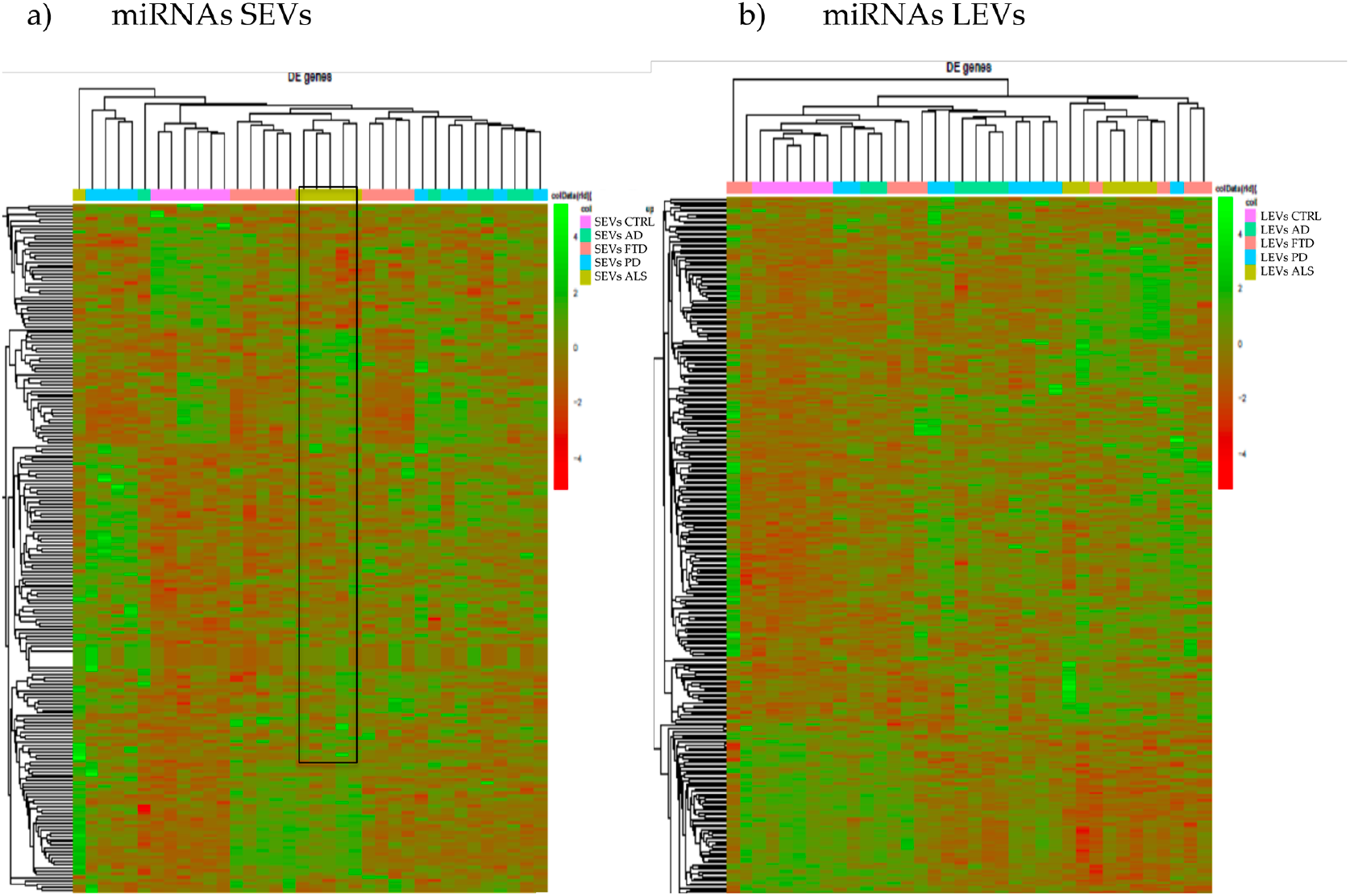
Heatmap of log-normalized miRNA expression counts for SEVs (a) and LEVs (b). All miRNAs identified as differentially expressed in at least one disease in the comparison of each disease to the control state were used to build the heatmap. Hierarchical clustering was applied to both X and Y axes. The black rectangle in figure a represent a specific signature of miRNAs with opposite expression to CTRs.

The PCA as well as the heatmap of the differential expressed miRNAs detected in LEVs did not provide a clear separation of the four diseases (Figure 3b, Figure 4b).

However, miRNAs in LEVs of ALS patients could be distinguished from CTRs and from the other three diseases, with some overlap with FTD.

In order to detect common miRNAs of the four diseases, we calculated the intersection of deregulated miRNAs compared to CTRs. We observed that 6 miRNAs were in common among the four diseases in SEVs (hsa-miR-133a-3, hsa-miR-543, hsa-miR-4451, hsa-miR-6889-5p, hsa-miR-4781-3p, hsa-miR-323b-3p) (Figure 5a) and 7 miRNAs (hsa-miR-1262, hsa-miR-3152-3p, hsa-miR-7856-5p, hsa-miR-365a-5p, hsa-miR-4433b-5p, hsa-miR-6068, hsa-miR-767-3p) in LEVs (Figure 5b). Enriched pathways found with MiRWalk included Ubiquitin mediated proteolysis, MAPK signaling pathway, Toll-like receptor signaling pathways for SEVs and Neurotrophin signaling pathway, MAPK signaling pathway, Glycosphingolipid biosynthesis, Ras signaling pathway for LEVs (Figure 5a and 5b).

**Figure 5.**
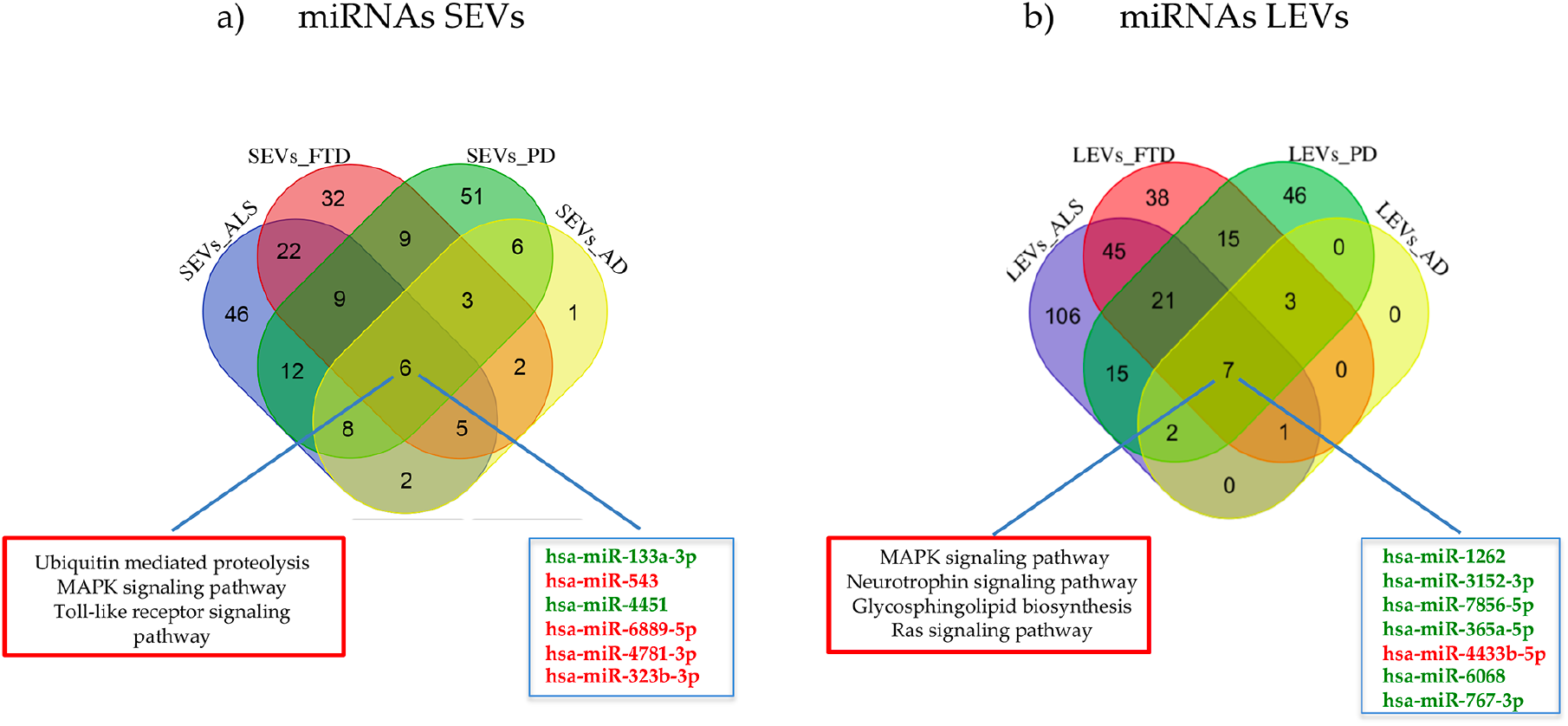
Venn diagram showing numbers of common and unique miRNA and RNA in SEVs (A) and LEVs (B) from plasma of AD, FTD, ALS and PD patients. Common miRNAs and pathways are listed. Differential mRNA expression analysis by edgeR (log2FC > 1, p-value<0.05).

### Coding and lncRNAs expression profiles and common pathways in SEVs and in LEVs of NDs

We analyzed differentially expressed mRNAs and lncRNAs in SEVs and LEVs of the four groups (Table 4, Table S2). As for miRNAs, mRNAs in SEVs showed that ALS patients are well divided from CTRs and from the other NDs except from FTD (Figure 6a), showing again a partial overlapping between ALS and FTD patients. In SEVs from AD and PD patients, no mRNAs and lncRNAs were found to be deregulated with |Log2FC| of 1.

**Figure 6.**
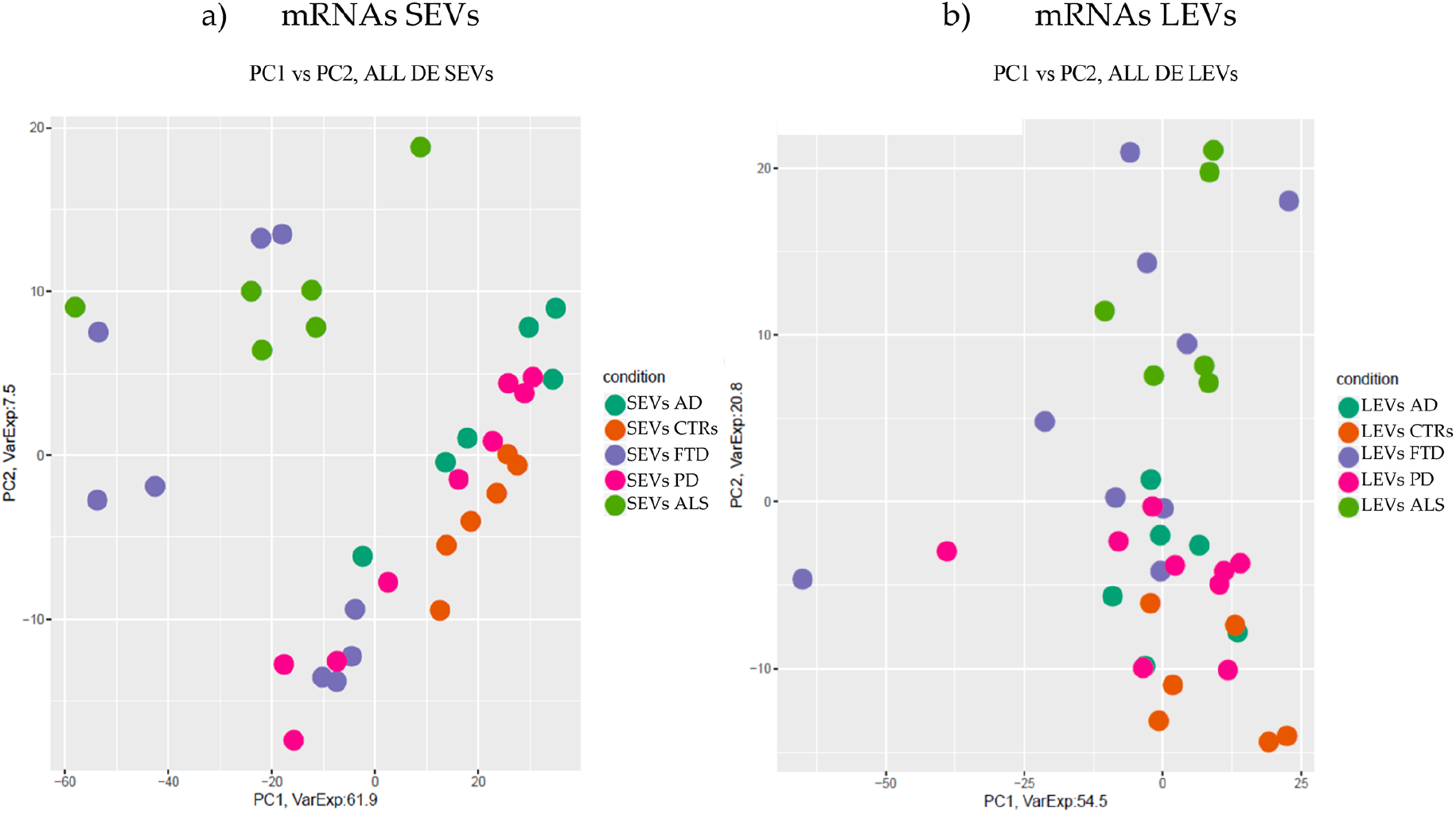
PCA of coding genes differentially expressed in SEVs (A) and LEVs (B) of AD, . FTD, ALS and PD patients and healthy controls (CTRs). PCA is performed using as predictors all the coding genes identified as differentially expressed in at least one disease in the comparison of each disease to the control state. Each dot represents a sample and each color represents a disease.

Considering only lncRNAs, both SEVs and LEVs showed a mixed scenario between patients and controls, without any specific characterization of AD, FTD . ALS or PD (Figure 7a and 7b).

**Figure 7.**
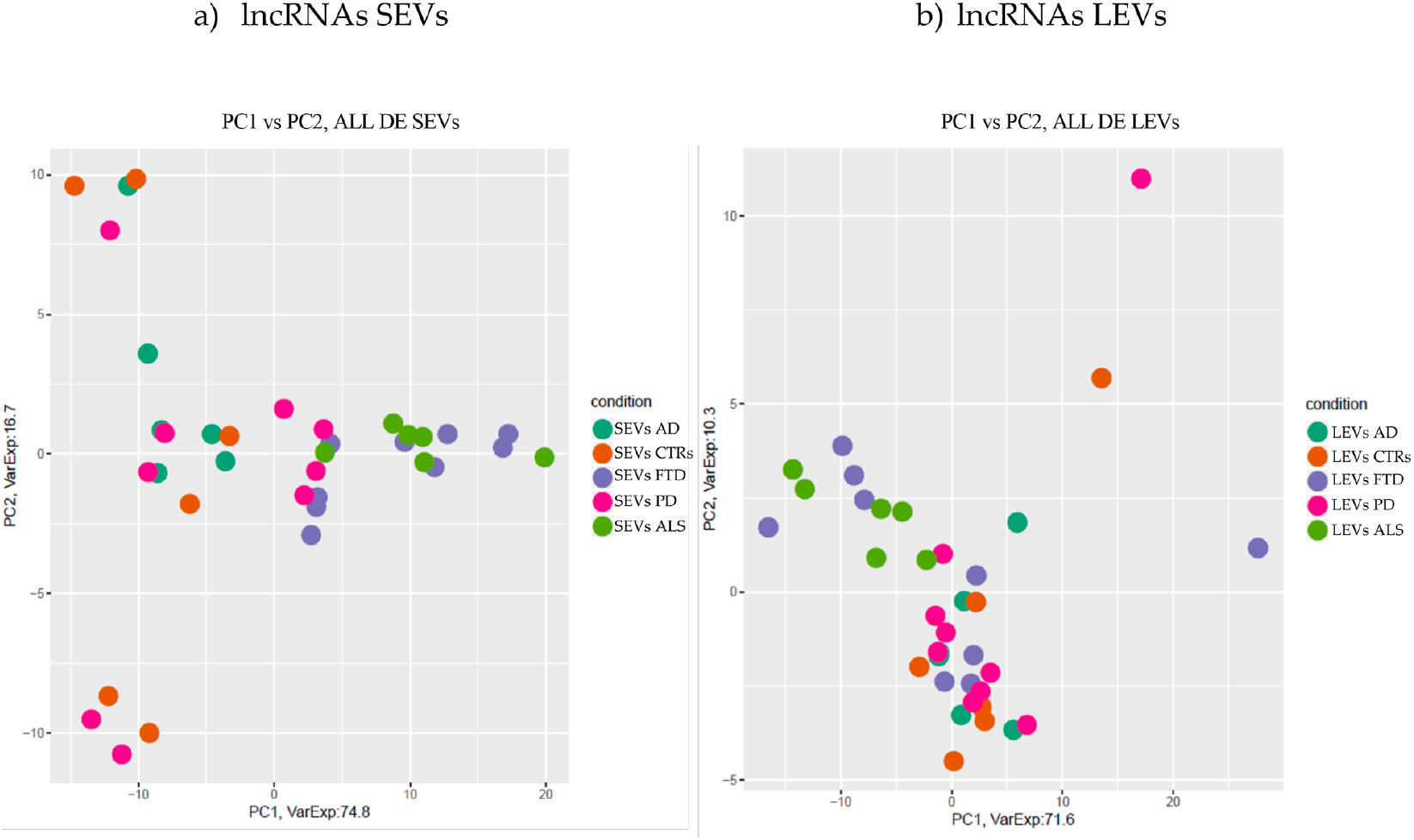
PCA of long non coding genes differentially expressed in SEVs (A) and LEVs (B) of AD, FTD, ALS, and PD patients and healthy controls (CTRs). PCA is performed using as predictors all the non-coding genes identified as differentially expressed in at least one disease in the comparison of each disease to the control state. Each dot represents a sample and each color represents a disease

In SEVs from FTD and ALS patients there were 235 genes in common and the common pathways of gene ontology biological processes (GO-BPs) were mRNA processing (p=6.133e-15), RNA splicing, via transesterification reactions with bulged adenosine as nucleophile and via spliceosome (Figure 8a). In LEVs there were only 2 genes in common among PD, FTD and ALS: MAP3K7CL, a kinase gene, and AP003068, 23, a transcript of an unknown protein (Figure 8b).

**Figure 8.**
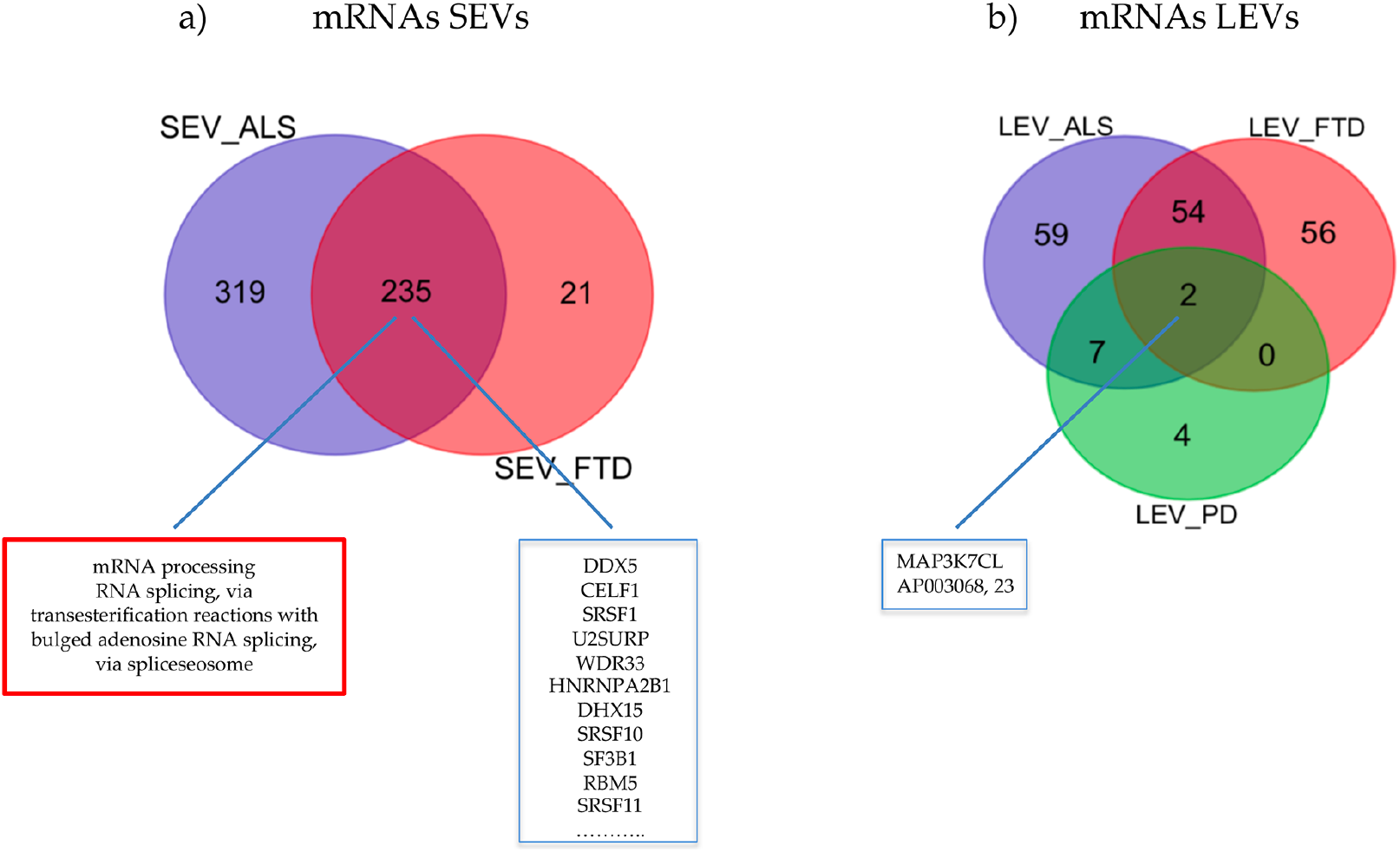
Venn diagram showing numbers of common and unique RNA in SEVs from plasma of ALS and FTD patients (A) and in LEVs from plasma of ALS, FTD and PD. Common miRNAs and pathways are listed. Differential mRNA expression analysis by DESeq2 (log2FC > 1, p-value<0.05).

### Specific miRNAs pathway analysis in SEVs and LEVs of NDs

#### 1. MiRWalk analysis and Ingenuity Pathway Analysis in SEVs of NDs

Enriched pathways targeted by the miRNAs that were differentially expressed between disease samples and CTRs were investigated by MiRWalk (Table S3).

Reactome and Gene Ontology (GO) analysis for DE miRNAs were run only on differentially expressed miRNAs compared to CTRs and not in common with the other NDs (Table S3). In SEVs from ALS patients Reactome analysis showed miRNAs involved in MECP2 (methyl CpG binding protein 2) expression and activity, Intracellular signaling by second messengers and negative regulators of DDX58/IFIH1. IFIH1 and DDX58 encode retinoic acid-inducible gene I (RIG-I), cytosolic pattern recognition receptors function in viral RNA detection initiating an innate immune response through independent pathways that promote interferon expression [48]. Ingenuity pathway analysis confirmed regulation of the immune system by highlighting how miR31-5p and miR615-5p control CXCL8, C-X-C Motif Chemokine Ligand 8 activated in viral infections and miR584-5p/miR148-5p control TMEM9, which enhances production of proinflammatory cytokines induced by TNF, IL1B, and TLR ligands [48]. GO-BPs found negative regulation of Ras protein signal transduction and very interestingly cell proliferation in forebrain.

In SEVs from FTD patients Reactome classified deregulated miRNAs in MyD88 cascade initiated on plasma membrane, diseases of signal transduction and axon guidance. GO-BPs classified deregulated miRNAs in SEVs from FTD in pathway release of cytochrome c from mitochondria, mitotic G1 DNA damage checkpoint and DNA damage response, signal transduction by p53 class mediator resulting in cell cycle arrest. IPA analysis (Figure 9b) confirmed involvement in MyD88 cascade (Adapter protein involved in the Toll-like receptor and IL-1 receptor signaling pathway in the innate immune response) and DNA damage response by regulation of miR146a-5p on RAD54, involved in homologous recombination and repair of DNA and on IL1F10 and TLR10 [49, 50] (Figure 9b).

**Figure 9.**
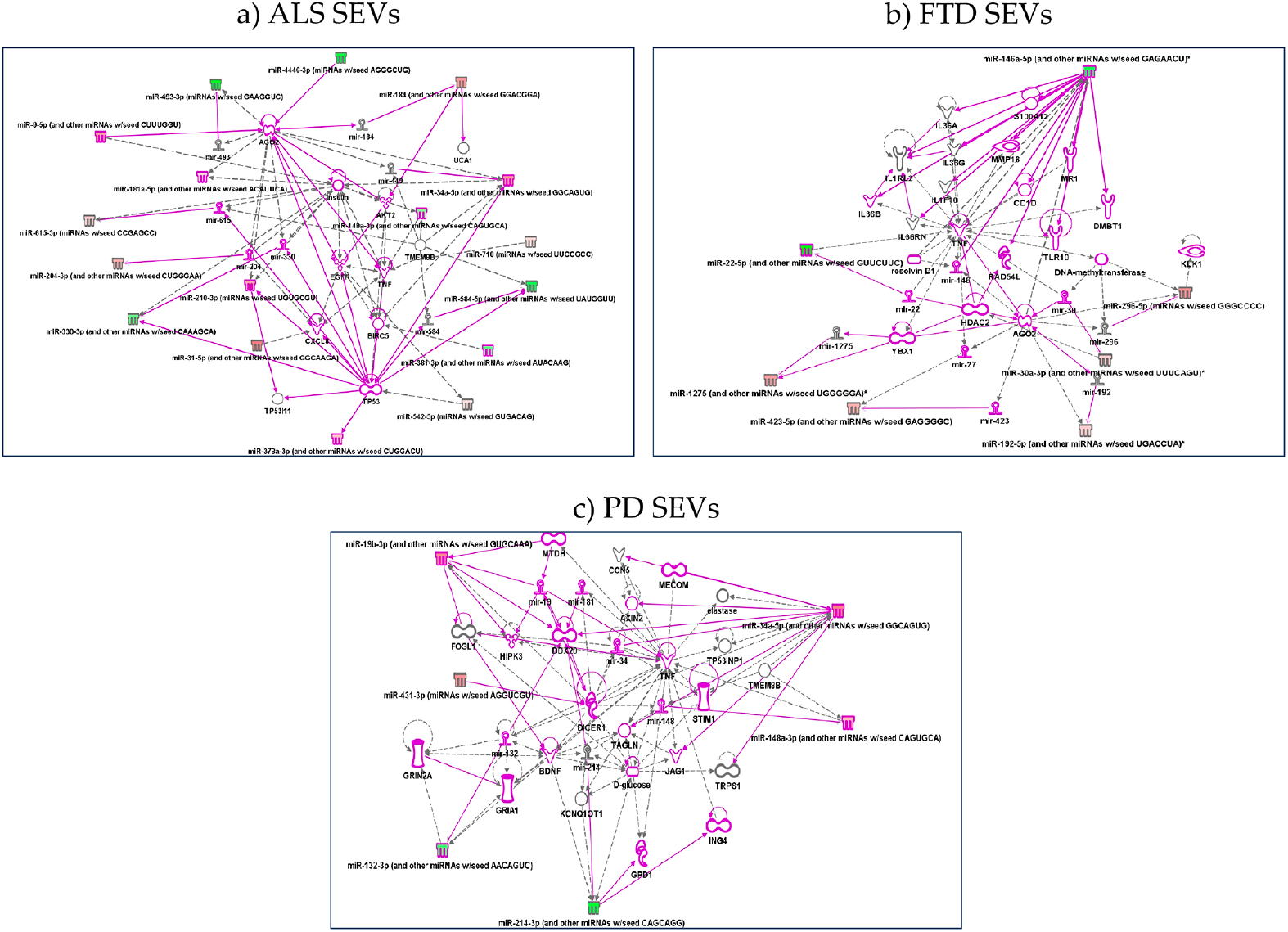
IPA networks among deregulated miRNAs in SEVs from ALS a), FTD (b), PD (c). For ALS miRNAs involvement in regulation of DDX58/IFIH1, through miR31-5p and miR615-5p (a); for FTD IL1F10 and TLR10 regulation by miR146a-5p (B); for PD, regulation of GRIA1, Glutamate receptors, by miR-132-3p, involvement of JAG1 and TNF regulated by miR-34 (c). Pink color indicates activation while green color indicate suppression. No pathway could be calculated for AD disease for the few miRNAs targets.

In SEVS from PD patients Reactome analysis identified signaling by TGF-beta family members, SLC-mediated transmembrane transport (regulation of GRIA1, subunit of Glutamate receptors, by miR-132-3p, Metabolism of lipids, MyD88 cascade initiated on plasma membrane (involvement of JAG1 and TNF regulated by miR-34-Sup IPA), while GeneOntology BP fibroblast migration, mRNA polyadenylation and positive regulation of gene silencing by miRNA (Figure 9c).

For AD, there were very few deregulated miRNAs to be classified in pathways.

#### 2. Up- and Down-regulated MiRWalk analysis in SEVs of NDs

Upregulated and downregulated miRNAs are listed in Table S4. In SEVs from ALS patients upregulated miRNAs could regulate MECP2, methyl CpG binding protein 2, expression and activity and RNA Polymerase III Transcription Initiation from Type 3 Promoter, while downregulated miRNAs regulate genes in post-translational protein modification and gene expression. In contrast to ALS, miRNAs only in SEVs from FTD patients are mainly downregulated compared to CTRs. These regulate genes in MyD88 cascade initiated on plasma membrane and in diseases of signal transduction. In SEVs from PD patients, the upregulated and downregulated miRNAs were not significantly associated with any biological pathways.

#### 3. MiRWalk analysis and Ingenuity Pathway Analysis in LEVs of NDs

Reactome underlined the role of miRNAs in LEVs: 1) from ALS patients in intracellular signaling by second messengers (DAG, cAMP, cGMP, IP3, Ca2+ and phosphatidylinositols), signaling by TGF-beta family members, MyD88 cascade initiated on plasma membrane and metabolism of lipids; 2) from FTD patients in gene expression; 3) from PD patients in intracellular signaling by second messengers, SLC-mediated transmembrane transport (solute carrier superfamily), some of which mediate neurotransmitter uptake in the CNS and peripheral nervous system (PNS) and metabolism of lipids.

Gene ontology analysis instead found pathways for LEVs: 1) from ALS patients regulation of G0 to G1 transition; 2) from FTD patients regulation of type I interferon-mediated signaling pathway and positive regulation of gene and posttranscriptional silencing; 3) from PD patients protein insertion into membrane, fibroblast apoptotic process, regulation of steroid biosynthetic process. Pathways are reported below in order of most significant p values and further details are shown in Table S3. IPA analysis confirmed the pathway found by both Reactome and Gene Ontology. 1) for LEVs from ALS regulation of p53 by mir-379, 378, 584-5p, 1207 emerged OR hsa-miR-199a-5p and hsa-miR-329a-5p regulate LCN2, lipocalin2, inducible factor secreted by reactive astrocytes in transgenic rats with neuronal expression of mutant human TDP-43 or RNA-binding protein FUS and that is selectively toxic to neurons [51] (Figure 10a); 2) for LEVs from FTD patients regulation of genes like TP53 by miR-296 and RNA Polymerase II by miR-615 as well as Cyclin E regulated miR-615-5p, that might be part of gene expression and regulation of apoptosis [52] (Figure 10b); 3) for LEVs from PD patients regulation of ADAM9 (ADAM Metallopeptidase Domain 9), important in mediating cell-cell and cell-matrix interactions, by upregulation of miR-291, miRNA which regulates cell proliferation and resolvin, a metabolic byproduct of omega-3 fatty acids, regulated by miR-302 [53,54] (Figure 10c).

**Figure 10.**
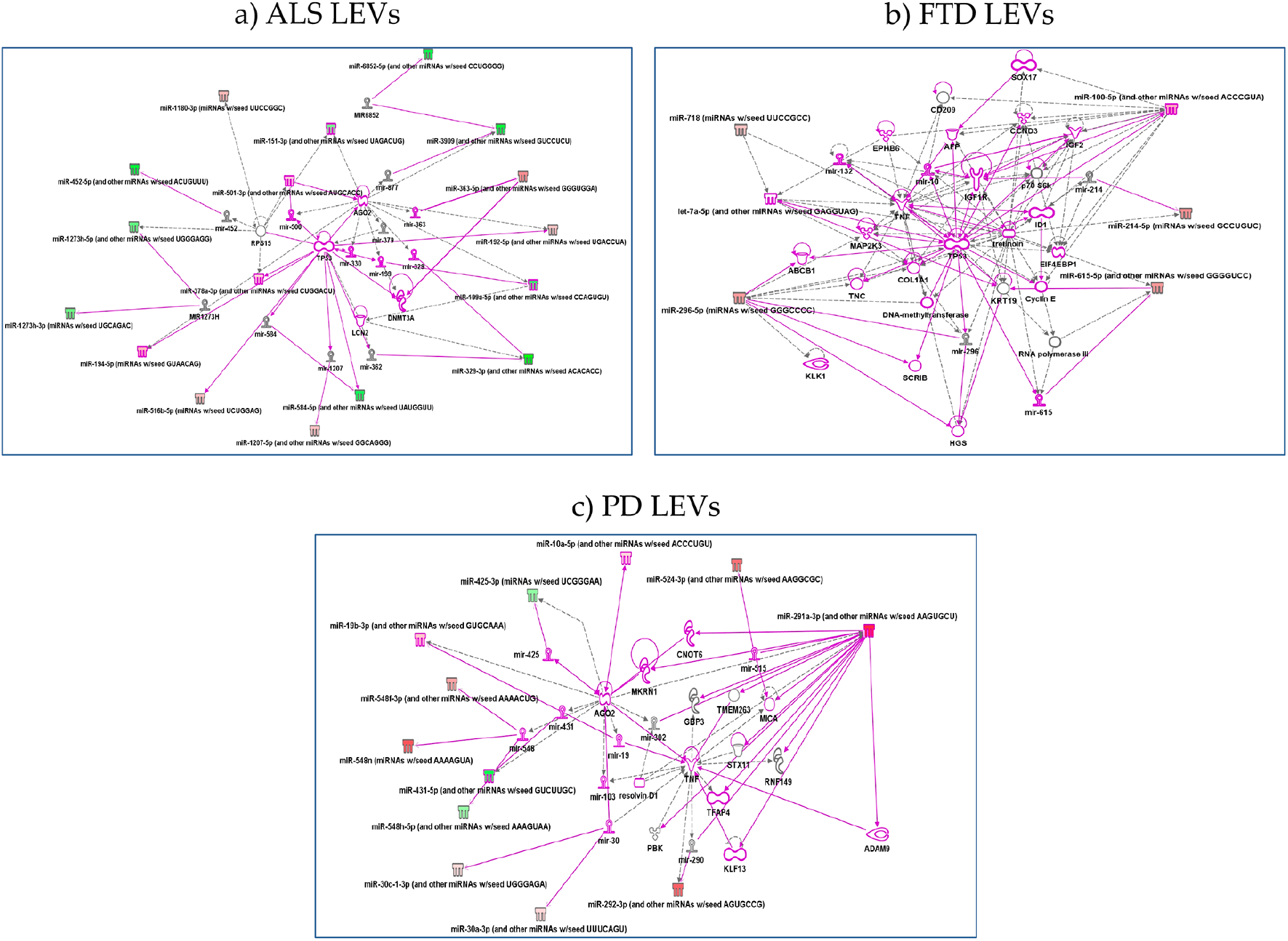
IPA networks among deregulated miRNAs in LEVs from ALS (A), FTD (B), PD (C). a) for ALS regulation of p53 by mir-379, 378, 584-5p, 1207 emerged or hsa-miR-199a-5p and hsa-miR-329a-5p regulate LCN2, lipocalin2); b) for FTD regulation of genes like TP53 by miR-296 and RNA Polymerase II by miR-615 as well as Cyclin E regulated miR-615-5p; c) for LEVs from PD patients regulation of ADAM9 by upregulation of miR-291. Pink color indicates activation while green color indicate suppression.

#### 4. Up- and Down-regulated <u>MiRWalk analysis in</u> LEVs of NDs

In LEVs from ALS patients, upregulated miRNAs belong to MyD88 cascade initiated on plasma membrane and SLC-mediated transmembrane transport (solute carrier superfamily), some of which mediate neurotransmitter uptake in the CNS and PNS, while downregulated miRNAs were related to intracellular signaling by second messengers (Reactome).

In LEVs from FTD patients, miRNAs regulating MECP2 expression and activity, MAPK signaling pathway and metabolism of lipids were upregulated.

In LEVs from PD patients, interestingly, most of the miRNAs are upregulated and the classes identified belonged to RAB geranylgeranylation, involved in trafficking of proteins in the endolysosomal system and in metabolism of lipids (Table S4).

For AD either the differentially expressed miRNAs were few, so it was not possible to calculate any pathway.

#### 5. Pathways analysis of mRNAs in SEVs and LEVs

EnrichR analysis (KEGG pathway and Gene Ontology (GO) analysis) for DEGs in FTD and ALS patients compared to healthy controls has been performed (Table S5). In ALS and FTD SEVs, GO BP analysis showed an important involvement of spliceosome and RNA metabolism as also reported above in section 3.2. The importance of transcription and RNA metabolism also emerged from KEGG analysis in SEVs and LEVs especially for ALS and for SEVs of FTD patients, showing an alteration of RNA degradation and transport pathways (Table S5). In LEVs of FTD patients, negative regulation of Ras protein signal transduction and positive regulation of neuron death were the most significant classes. In LEVs for PD only 13 genes were deregulated compared to CTRs and pathways like the biosynthesis of unsaturated fatty acids were statistically significant (Table S5).

We also looked at DEGs compared to CTRs and not in common to the other NDs. GO BP identified 1) in LEVs from ALS, positive regulation of transmembrane transport, positive regulation of leukocyte chemotaxis; 2) in LEVs from FTD cellular response to molecule of bacterial origin; 3) in LEVs from PD positive regulation of mitochondrial membrane permeability involved in apoptotic process, mitochondrial outer membrane permeabilization; 4) in SEVs from ALS regulation of transcription from RNA polymerase II promoter, regulation of transcription, DNA-templated, negative regulation of gene expression; 5) in SEVs from FTD patients regulation of cellular catabolic process, phosphatidylinositol metabolic process, transcription initiation from RNA polymerase III promote (Table S6).

## Discussion

Identifying robust biomarkers is essential for early diagnosis of NDs. Extracellular vesicles (SEVs and LEVs), transported in blood, might play this role. In this study, we have analyzed SEVs and LEVs cargo in order to detect RNAs acting as novel, easily accessible biomarkers for AD, PD, ALS and FTD.

We first compared miRNA and mRNA expression profiles of SEVs and LEVs in the same disease. We found a variable range of overlap between LEVs and SEVs: for miRNAs in SEVs the percentage was between 18.2-61.5% and in LEVs 25.2-46.2% and for mRNAs 8.4 and 17.1% for SEVs and 35.5 and 34.2 for ALS and FTD patients (Table 5). Although there is some overlap between the two types of EVs, there is a significant difference that may justify, as we already described for dimension, protein and lipid loading [35,40,55], the different functions of LEVs and SEVs in plasma of ND patients. Conley et al. characterized protein coding transcripts in SEVs and LEVs from breast cancer patients by RNA-Seq and identified a small fraction of transcripts that were expressed at significantly different levels in large oncosomes and exosomes, suggesting they may mediate specialized functions [56].

Regarding common deregulated miRNAs in LEVs and SEVs in ALS, our data showed an important deregulation in a small group of specific miRNAs already described in the literature (hsa-miR-206, hsa-miR-205-5p, miR-1-3p, hsa-miR-205-5p, hsa-miR-200b-3p, hsa-miR-200c-3p, hsa-miR-6888-3p, hsa-miR-31-5p, hsa-miR-141-3p, hsa-miR-210-3p). MiR-206 is the main described miRNA associated to ALS [57,58]. MiR-1 and miR-206 has already been described to be deregulated in ALS patients [57,59], while in previous studies miR-141 and miR-200 were reported as related to ALS since they bind a sequence in FUS promoter and these mRNAs are linked by a feed-forward regulatory loop where FUS upregulates miR-141/200a, which in turn impact FUS protein synthesis [60,61]. Also, miR-210 is already known to be up-regulated in neurodegenerative diseases [62]. To our knowledge, remaining miRNAs deregulation was never reported as associated to ALS and may represent novel biomarkers specific for this disease.

We then analyzed deregulated miRNAs and mRNAs among the four NDs by the principal component analysis (PCA). We found that deregulated miRNAs cargo of the four NDs was different from CTRs in particular in SEVs, while in in LEVs the only group that did not overlap with CTRs was ALS, and so it was for PCA related to mRNAs in LEVs and SEVs. Specifically, in PCA of miRNAs of SEVs the three groups, CTRs, ALS and AD, were well separated suggesting that miRNAs cargo of SEVs might be bound to ND phenotypes. On the other hand, miRNAs split the group of PD patients in two, one overlapped with AD and the other with FTD patients. The overlap of miRNA signature cannot be justified by similar clinical phenotypes among the groups of patients: of the five PD patients which overlap with AD patients, only one displayed cognitive deficit and the other one had Lewy Body Dementia. Also, FTD patients showed two subgroups, one overlapping with PD and the other with ALS. Multiple studies [63,64,65] already demonstrated that the observed overlap between FTD and ALS is due to common mechanisms contributing to the onset and development of the disease. Also in this case there is no correlation with the clinical history. Parkinsonism is found in approximately 20–30% of patients in FTLD, in particular it is frequently observed in familial FTD, with mutations linked to microtubule associated protein Tau (MAPT), progranulin (GRN or PGRN), and chromosome 9 open reading frame 72 (C9ORF72) repeat expansion [66]. In our cohort, only one FTD patient presented parkinsonism, which, however, clustered in the ALS group and did not present any mutation in the canonical genes associated to familial FTD. All FTD patients showed variable amounts of Tau, ßamyloid 1-42 and only two patients presented ALS-FTD disease.

As shown in Table 3 and Table 4, EVs were enriched with a greater number of DE miRNAs compared to mRNAs. In diseases like ALS and FTD, SEVs were enriched also with a greater number of deregulated mRNAs, while diseases like PD and in particular AD have a minor number of DE mRNA compared to CTRs. It is also clear that SEVs are more enriched in deregulated miRNAs compared to LEVs. This is in agreement with the literature demonstrating that SEVs contain primarily small RNA [67]. In fact, in the panel of neurodegeneration, the data about RNA metabolism and AD are few [68], suggesting that the RNA regulation does not have a fundamental role in the mechanism of the pathology.

Common miRNAs in the four NDs were different between the two groups of EVs. Enriched pathways of common miRNAs found with MiRWalk showed pathway like Ubiquitin mediated proteolysis, MAPK signaling pathway, Toll-like receptor signaling pathways for SEVs and TGF-beta signaling pathway, Neurotrophin signaling pathway, MAPK signaling pathway, Glycosphingolipid biosynthesis, Ras signaling pathway for LEVs. It is interesting how deregulated common miRNAs both in LEVs and SEVs are involved in signal transduction. Although recent reports have implicated EVs in intercellular signaling [69], their influence in modulating signaling pathways in the target cells is not fully clear.

For the common enriched pathway of SEVs in the four NDs, it is known that many NDs are related to inflammation, which can increase cell injury and cause neuronal death. Toll like receptors (TLR) are innate immune receptors that, when activated, can induce the downstream signal molecules through the MyD88-dependent and TRIF signal adaptor proteins, which activate downstream kinases including IkB kinases and MAP kinases. In general, TLRs are expressed in the CNS, in neurons (TLR3, 4, 7, and 9), in human oligodendrocytes (TLR2), in human astrocytes (TLR3-4), and in human microglia (TLR1-4). Activation of both endosomal and plasma membrane receptors like TLRs can activate microglia and control the evolution of neurodegenerative processes [70].

Ubiquitin mediated proteolysis is the process of degradation of a protein via the ubiquitin system [71] and it is one of the common pathways of SEVs from the four NDs that we found. NDs are characterized by intraneuronal inclusions containing ubiquitynated filamentous protein aggregates, given by loss of function or mutations in enzymes of the ubiquitin conjugation/deconjugation pathway [72]. If there is an impairment of the ubiquitin mediated proteolysis, SEVs, which originate in the endocytic pathway, (differently from LEVs, which are shed from the budding of the cell membrane) might be the affected key part of the machinery as already suggested. In fact, the incorporation of ubiquitinated proteins into intraluminal vesicles (ILVs) is controlled through the Endosomal Sorting Complexes Required for Transport, ESCRT complex, key part of SEVs [73].

For LEVs, Neurotrophin signaling pathway, also called nerve growth factor (NGF) family members pathway, has multiple functions in both developing and mature neurons and it is connected to the downstream MAPK and Ras signaling pathway. On activation by BDNF, trkB initiates intracellular signaling through Shc and PLCγ binding sites. The Shc binding site plays major roles in neuronal survival and axonal outgrowth [74]. Another common deregulated pathway is glycosphingolipid biosynthesis pathway. Increasing evidence underlines the activation of ceramide-dependent pro-apoptotic signaling and reduction of neuroprotective S1P in neurodegeneration course. One hypothesis is the link between altered ceramide/S1P and the production, secretion, and aggregation of pathological proteins in NDs. Sphingolipids regulate EVs and the spread or release of neurotoxic proteins and/or regulatory miRNAs between brain cells [75].

The expression profile of some miRNAs has already been described in the brain or in blood of some NDs, but with an opposite regulatory pattern to the one found in EVs of our study.

For example, miR-133b expression is downregulated in the midbrain of PD patients and in an animal model of PD, while in SEVs is upregulated [76]. miR-4781–3p was instead found upregulated in blood of AD patients by RNAseq, while in SEVs of the four NDs, this miRNA is downregulated in SEVs [77]. miR-323–3p associated with inflammatory responses, has been proposed as a target for therapy in AD and it is found downregulated in SEVs [78]. Some of these miRNAs have not been found in neurodegeneration, and some only in one of the NDs studied in this work. Further studies are needed in this regard and specific miRNAs and pathways will be discussed in future works for each disease in future works.

In this report, we have also investigated the regulation of coding and lncRNAs. Coding RNAs showed a similar picture to miRNAs pattern for ALS disease. As for miRNAs, coding RNAs in SEVs of ALS better separate from other disease compared to CTRs. Notably, in the list of deregulated mRNAs in ALS patients, we have found 3 RNA Binding Motif Protein, a protein family already associated to ALS [79]. In contrast, deregulated lncRNAs did not cluster apart in the different diseases and, in AD and PD patients, no lncRNAs were found to be deregulated. Of the common genes between ALS and FTD, they were mainly classified as belonging to splicing, a mechanism largely described in those two NDs [80]. Further studies are needed on extended cohorts of patients with different stages of the disease in order to understand if the deregulation of these RNAs may be associated to a specific clinical window (disease onset and outcome) and progression of the disease.

## Conclusions

We found different deregulation of miRNAs and mRNAs between SEVs and LEVs from plasma of patients in four NDs.

Deregulated cargo in SEVs may be a starting point for a specific signature for ALS disease, paving the way for future studies on a specific small group of miRNAs that may become peripheral ALS biomarkers.

We found a common signature of miRNAs in SEVs and LEVs among the four NDs and those miRNAs are involved in pathways already known in neurodegeneration.

The novelty is that different EVs are involved in different pathways and this might give a specificity to the role of SEVs and LEVs in the spreading/protection of the disease.

## Abbreviations

AD: Alzheimer’s Disease
ALS: Amyotrophic Lateral Sclerosis
CTRs: healthy controls
DE miRNAs: differentially expressed miRNAs
DE mRNA: differentially expressed mRNAs
ESCRT: Endosomal Sorting Complexes Required for Transport
EVs: extracellular vesicles
EXOs: exosomes
FTD: Frontotemporal Dementia
GWAS: Genome-wide association studies
ILVs: intraluminal vesicles
IPA: Ingenuity pathway analysis
LBs: Lewy bodies
LEVs: large extracellular vesicles
lncRNA: long non coding RNA
miRNA: microRNA
MVs: microvesicles
NDs: neurodegenerative diseases
PCA: principal component analysis
PD: Parkinson’s Disease
RBPs: RNA-binding proteins
SEVs: small extracellular vesicles
SN: substantia nigra
SNPs: single nucleotide polymorphisms
TLRs: Toll like receptors

## Declarations

### Ethical Approval and Consent to participate

Subjects participating in the study signed, before being enrolled, an informed consent form approved by the Ethical Committee of IRCCS Mondino Foundation, Pavia, Italy (for ALS patients Protocol n°-20180034329; for PD patients Protocol n°20170001758; for AD patients Protocol n°20170016071; for FTD patients Protocol n°20180049077).

### Consent for publication

Not applicable

### Availability of data and materials

The datasets supporting the conclusions of this article are included within the article and its additional files.

### Authors’ contributions

Writing—review and editing, S.G., D.S. and S.Z.; D.S., M.A. and M.G. performed the experiments. R.C., S.Z., and M.O. performed bioinformatic analysis. S.B., M.A., A.C., B.M., G.P., L.D., R.Z., M.C.R., M.C. participated to patients and controls recruitment. M.A., D.S., O.P. and C.C. set up the experimental plan. O.P., R.C. and C.C. review. R.C. and C.C. supervision. All authors reviewed and accepted the final version of this manuscript.

## Acknowledgements

We kindly thank Prof. Fabio Corsi and Mr. Raffaele Allevi (Dipartimento di Scienze Biomediche e Cliniche “L. Sacco”) for TEM images and analysis. We thank patients and family for contributing to this study.

## Additional files

**Figure S1: LEVs and SEVs characterization.** a) Nanosight profile of LEVs and SEVs from plasma of a CTR, AD, PD, ALS, FTD patients; b and c) Representative images obtained by transmission electron microscopy (TEM) of LEVs and SEVs from plasma (Scale bar: 100 nm, 50 nm). d) Western Blot of LEVs and SEVs markers in LEVs and SEVs samples from one CTR, an ALS, AD, PD, FTD patients showed the presence of Annexin V only in LEVs pellet and Alix in SEVs fraction.

**Table S1. Differentially expressed miRNAs in ALS, FTD, PD and AD groups respect to healthy controls.** miRNA ID, measured log2FC and p-value are reported for each transcript.

**Table S2. Differentially expressed RNAs in ALS, FTD, PD and AD groups respect to healthy controls.** Transcript ID, gene name, gene type, gene status, measured log2FC and false discovery rate (FDR) are reported for each transcript. Only transcripts with with |log2(disease sample/healthy control)|≥1 and a FDR ≤ 0.1 are shown.

**Table S3. miRWalk specific miRNAs pathways in ALS, FTD, PD and AD groups.** Deregulated miRNAs specific of each disease were analysed with miRWalk web tool. Reactome Analysis and Gene Ontology Biological Processes with a p value< 0.05 are listed.

**Table S4. miRWalk specific miRNAs pathways divided in up and down-regulated miRNAs in ALS, FTD, PD and AD groups.** Deregulated miRNAs specific of each disease were analysed with miRWalk web tool. Reactome Analysis and Gene Ontology Biological Processes with a p value< 0.05 are listed.

**Table S5. mRNAs pathways in ALS, FTD, PD and AD groups.** EnrichR web tool was used to calculate enriched pathways. KEGG and Gene Ontology Biological Processes with a p value< 0.05 are listed

**Table S6. Specific mRNAs pathways in ALS, FTD, PD and AD groups.**Deregulated mRNAs specific of each disease were analysed with EnrichR web tool. KEGG and Gene Ontology Biological Processes with a p value< 0.05 are listed

