## Supplementary material for "Different RNA profiles in plasma derived small and large extracellular vesicles of Neurodegenerative diseases patients": Figure S1

### Slide 1
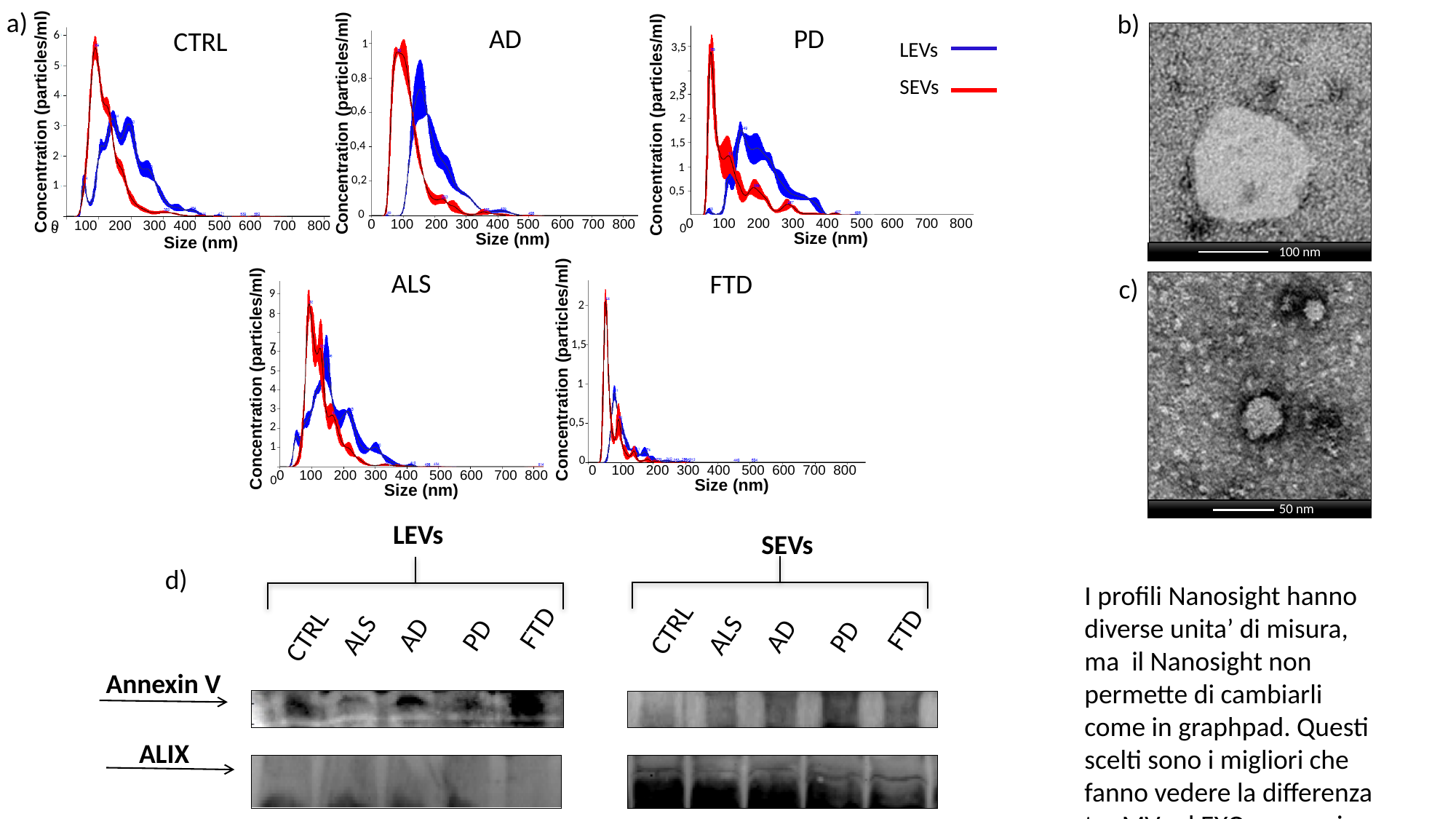

a)
CTRL
6
5
Concentration (particles/ml)
4
3
2
1
 0
0 100 200 300 400 500 600 700 800
Size (nm)
b)
AD
PD
3,5
 3
2,5
2
1,5
1
0,5
 0
0 100 200 300 400 500 600 700 800
Size (nm)
1
LEVs
0,8
SEVs
Concentration (particles/ml)
Concentration (particles/ml)
0,6
0,4
0,2
FTD
2
Concentration (particles/ml)
1,5
1
0,5
0
0 100 200 300 400 500 600 700 800
Size (nm)
0
ALS
9
8
 7
6
Concentration (particles/ml)
5
4
3
2
1
 0
Size (nm)
0 100 200 300 400 500 600 700 800
0 100 200 300 400 500 600 700 800
Size (nm)
100 nm
c)
50 nm
LEVs
SEVs
CTRL
AD
AD
FTD
CTRL
FTD
PD
PD
ALS
ALS
Annexin V
ALIX
d)
I profili Nanosight hanno diverse unita’ di misura, ma il Nanosight non permette di cambiarli come in graphpad. Questi scelti sono i migliori che fanno vedere la differenza tra MV ed EXO per ogni malattia e sono insicativi di un solo paziente come abbiamo fatto per gli altri 3 papers. Altrimenti
